# K18-hACE2 mice develop respiratory disease resembling severe COVID-19

**DOI:** 10.1101/2020.08.11.246314

**Authors:** Claude Kwe Yinda, Julia R. Port, Trenton Bushmaker, Irene Offei Owusu, Victoria A. Avanzato, Robert J. Fischer, Jonathan E. Schulz, Myndi G. Holbrook, Madison J. Hebner, Rebecca Rosenke, Tina Thomas, Andrea Marzi, Sonja M. Best, Emmie de Wit, Carl Shaia, Neeltje van Doremalen, Vincent J. Munster

## Abstract

SARS-CoV-2 emerged in late 2019 and resulted in the ongoing COVID-19 pandemic. Several animal models have been rapidly developed that recapitulate the asymptomatic to moderate disease spectrum. Now, there is a direct need for additional small animal models to study the pathogenesis of severe COVID-19 and for fast-tracked medical countermeasure development. Here, we show that transgenic mice expressing the human SARS-CoV-2 receptor (angiotensin-converting enzyme 2 [hACE2]) under a cytokeratin 18 promoter (K18) are susceptible to SARS-CoV-2 and that infection resulted in a dose-dependent lethal disease course. After inoculation with either 10^4^ TCID_50_ or 10^5^ TCID_50_, the SARS-CoV-2 infection resulted in rapid weight loss in both groups and uniform lethality in the 10^5^ TCID_50_ group. High levels of viral RNA shedding were observed from the upper and lower respiratory tract and intermittent shedding was observed from the intestinal tract. Inoculation with SARS-CoV-2 resulted in upper and lower respiratory tract infection with high infectious virus titers in nasal turbinates, trachea and lungs. The observed interstitial pneumonia and pulmonary pathology, with SARS-CoV-2 replication evident in pneumocytes, were similar to that reported in severe cases of COVID-19. SARS-CoV-2 infection resulted in macrophage and lymphocyte infiltration in the lungs and upregulation of Th1 and proinflammatory cytokines/chemokines. Extrapulmonary replication of SARS-CoV-2 was observed in the cerebral cortex and hippocampus of several animals at 7 DPI but not at 3 DPI. The rapid inflammatory response and observed pathology bears resemblance to COVID-19. Taken together, this suggests that this mouse model can be useful for studies of pathogenesis and medical countermeasure development.

**Authors Summary:** The disease manifestation of COVID-19 in humans range from asymptomatic to severe. While several mild to moderate disease models have been developed, there is still a need for animal models that recapitulate the severe and fatal progression observed in a subset of patients. Here, we show that humanized transgenic mice developed dose-dependent disease when inoculated with SARS-CoV-2, the etiological agent of COVID-19. The mice developed upper and lower respiratory tract infection, with virus replication also in the brain after day 3 post inoculation. The pathological and immunological diseases manifestation observed in these mice bears resemblance to human COVID-19, suggesting increased usefulness of this model for elucidating COVID-19 pathogenesis further and testing of countermeasures, both of which are urgently needed.

## Introduction

Severe acute respiratory syndrome coronavirus-2 (SARS-CoV-2) emerged in Hubai province in mainland China in December 2019, and is the etiological agent of coronavirus disease (COVID)-19 (1). SARS-CoV-2 can cause asymptomatic to severe lower respiratory tract infections in humans, with early clinical signs including fever, cough and dyspnea (2, 3). Progression to severe disease may be marked by acute respiratory distress syndrome (ARDS), with pulmonary edema, bilateral diffuse alveolar damage and hyaline membrane formation (4-6). Although primarily a respiratory tract infection, extra-respiratory replication of SARS-CoV-2 has been observed in kidney, heart, liver and brain in fatal cases (7-9). Several experimental animal models for SARS-CoV-2 infection have been developed, including hamsters (10) ferrets (11) and non-human primate models (12-15). SARS-CoV-2 pathogenicity within these animal models ranges only from mild to moderate (10-15). Additional small animal models that recapitulate more severe disease phenotypes and lethal outcome are urgently needed for the rapid pre-clinical development of medical countermeasures. Although the SARS-CoV-2 spike glycoprotein is able to utilize hamster angiotensin-converting enzyme 2 (ACE2) as the receptor of cell entry (10, 16), lack of species-specific reagents limit the usability of this model. As SARS-CoV-2 is unable to effectively utilize murine (m)ACE2 (17, 18), several models are currently under development to overcome this species barrier using a variety of strategies including transiently expressed human (h)ACE2, CRISPR/Cas9 modified mACE2, exogenous delivery of hACE2 with a replication-deficient viral vector and mouse-adapted SARS-CoV-2 (19-23).

K18-hACE2 transgenic mice were originally developed as a small animal model for lethal SARS-CoV infection. Expression of hACE2 is driven by a cytokeratin promoter in the airway epithelial cells as well as in epithelia of other internal organs, including the liver, kidney, gastrointestinal tract and brain. Infection with SARS-CoV led to severe interstitial pneumonia and death of the animals by day 7 post inoculation (20). Here, we assess the susceptibility of K18-hACE2 transgenic mice as a model of severe COVID-19.

## Results

### Disease manifestation in SARS-CoV-2-inoculated K18-hACE2 mice

First, we determined the disease progression after SARS-CoV-2 inoculation. Two groups of 4-6 week-old K18-hACE2 transgenic male and female mice (15 each) were intranasally inoculated with 10^4^ (low dose group) and 10^5^ (high dose group) TCID_50_ SARS-CoV-2, respectively. In addition, one control group of two mice was intranasally inoculated with 10^5^ TCID_50_ γ-irradiated SARS-CoV-2.

Irrespective of SARS-CoV-2 inoculation dose, mice uniformly started losing weight at 2 days post inoculation (DPI) (Fig 1a), with a significantly higher weight loss observed in the low dose group, suggesting a dose-response relationship, (p = 0.02, Wilcoxon matched-pairs rank test). No difference in weight loss between male and female animals within the same dose group was detected (S1a Fig). In addition to weight loss, lethargy, ruffled fur, hunched posture and labored breathing were observed throughout the course of infection in each animal. Mice were monitored for signs of neurological disease (circling, rolling, hyperexcitability, convulsions, tremors, weakness, or flaccid paralysis of hind legs), and no neurological symptoms were observed in any of the animals. Within the high dose group all animals reached euthanasia criteria by 7 DPI, however, in the low dose group five out of six animals reached euthanasia criteria 5-9 DPI and one animal recovered (Fig 1b). Although, no sex-dependent differences in survival were observed between male and female mice, the animal size used in this study was too small to draw major conclusions (S1b Fig). The control animals inoculated with γ-irradiated SARS-CoV-2 did not lose weight and remained free of disease symptoms.

**Fig 1.**
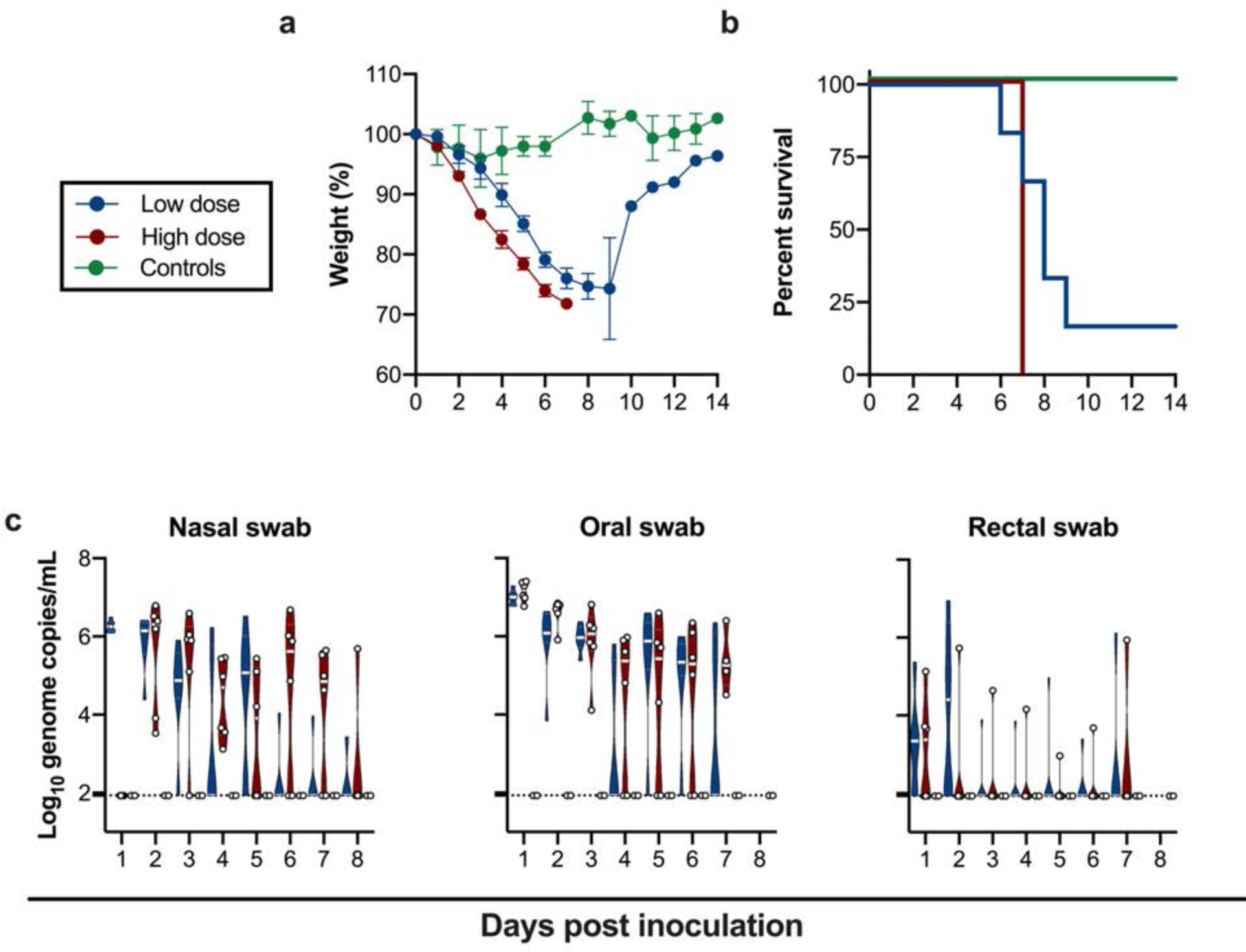
Inoculation of K18-hACE2 mice results in lethal infection and virus shedding. **a**. Relative weight loss in mice after SARS-CoV-2 inoculation. The lines represent mean ± SEM. **b**. Survival curves of mice inoculated with 10^4^ or 10^5^ TCID_50_ SARS-CoV-2, or 10^5^ γ-irradiated SARS-CoV-2. **c**. Violin plot of viral load in nasal, oropharyncheal and rectal swabs with geometric mean as centre. Viral RNA was quantified using RT-qPCR in nasal, oropharyncheal and rectal swabs, bar at geometric mean. Blue: 10^4^ TCID_50_ (low dose animals, n = 6); red: 10^5^ TCID_50_ (high dose animals, n = 6); green: 10^5^ TCID_50_ γ-irradiated (control animals, n = 2); dotted line = limit of detection.

### Viral shedding in SARS-CoV-2-inoculated K18-hACE mice

To gain an understanding of dose-dependent virus shedding patterns of SARS-CoV-2 in infected K18-hACE2 mice, daily nasal, oropharyngeal and rectal swabs were obtained until 11 DPI. Viral RNA was detected in all three. SARS-CoV-2 shedding from the respiratory tract was observed in all inoculated animals. Viral load in oropharyngeal and nasal swabs reached up to ∼10^6^ and ∼10^7^ copies/mL, respectively, and viral RNA could be detected up to 7 and 8 DPI. Rectal shedding was observed in both inoculated groups, but not in all animals, and was lower compared to respiratory shedding. Importantly, no viral RNA could be detected in swabs obtained from control mice inoculated with γ-irradiated SARS-CoV-2, suggesting viral RNA detected as early as 1 DPI was directly associated with active virus replication and did not originate from inoculum (Fig 1c). No sex-dependent differences in shedding pattern were seen (S1c Fig).

### Tissue tropism of SARS-CoV-2-inoculated K18-hACE mice

We next assessed tissue tropism and viral replication of SARS-CoV-2 in K18-hACE2 mice (Fig 2a). Viral genomic RNA was detected in almost all tissues; however, no viremia was observed. At 3 and 7 DPI, the highest viral load was found in lung tissue (∼10^10^ genome copies/g). Viral RNA in brain tissue was increased at 7 DPI compared to 3 DPI (from ∼10^5^ to 10^10^ genome copies/g) (Fig 2a). When assessing infectious virus, at 3 DPI, it was only detected in respiratory tract tissues, with high infectious titers observed in nasal epithelium and lungs in both the low dose and high dose groups. At 7 DPI, infectious virus was detected in respiratory tract as well as brain tissue (Figs 2b). Together, these data suggest that either SARS-CoV-2 is initially exclusively targets the respiratory tract in K18-hACE2 mice with secondary central nervous system (CNS) involvement or the virus replicates slower in the brain and only detected after 3 DPI.

**Fig 2.**
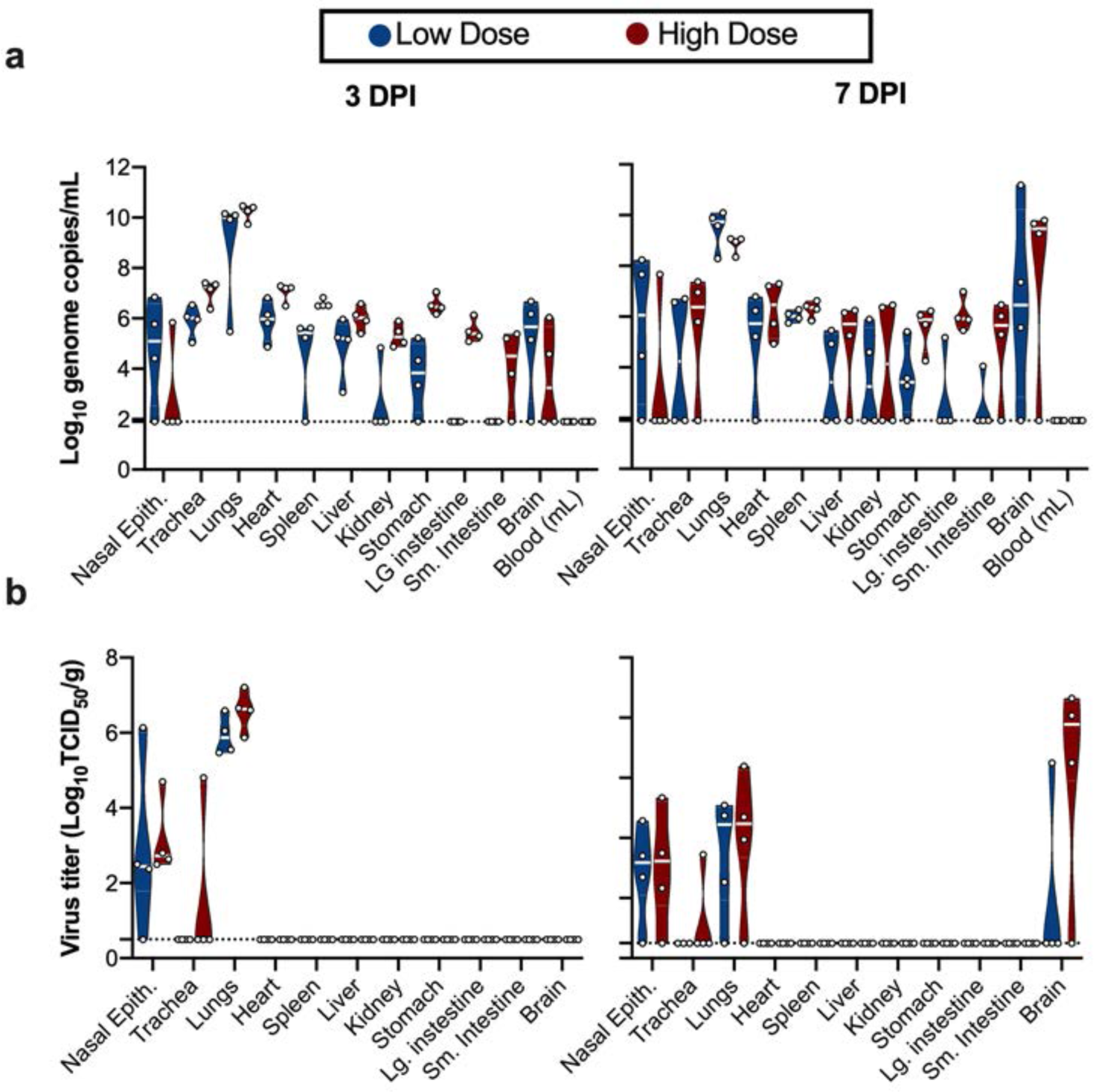
SARS-CoV-2 tissue tropism in K18-hACE mice. **a.** Violin plot of viral load in tissues quantified by UpE RT-qPCR with geometric mean as center. **b.** Violin plot of infectious SARS-CoV-2 titers in tissues, with geometric mean as centre. Blue: 10^4^ TCID_50_ (low dose animals, n = 6); red: 10^5^ TCID_50_ (high dose animals, n = 6); green: 10^5^ TCID_50_ γ-irradiated (control animals, n = 2); dotted line = limit of detection.

### Histological changes and viral antigen distribution in SARS-CoV-2-inoculated K18-hACE mice

On 3 and 7 DPI, four animals from each group were euthanized and necropsies were performed. On both days, gross lung lesions were observed in all animals with up to 80% of the lungs affected by 7 DPI. Histologically, all animals developed pulmonary pathology after inoculation with SARS-CoV-2. Lungs showed interstitial pneumonia at 3 DPI characterized by a generalized perivascular infiltration of inflammatory cells including neutrophils, macrophages and lymphocytes; alveolar septal thickening, and distinctive vascular system injury (Fig 3a-3c). At 7 DPI, mice developed pulmonary pathology consisting of multifocal interstitial pneumonia characterized by type II pneumocyte hyperplasia, septal, alveolar and perivascular inflammation comprised of lymphocytes, macrophages and neutrophils, variable amounts of alveolar fibrin and edema, frequent syncytial cells and single cell necrosis. Terminal bronchioles were similarly affected and in the most severely affected areas fibrin and necrosis occluded the lumen (Fig 3e-3g). Immunohistochemistry (IHC) demonstrated viral antigen in pneumocytes and macrophages of tissues on both 3 and 7 DPI (Fig 3d-3h).

**Fig 3.**
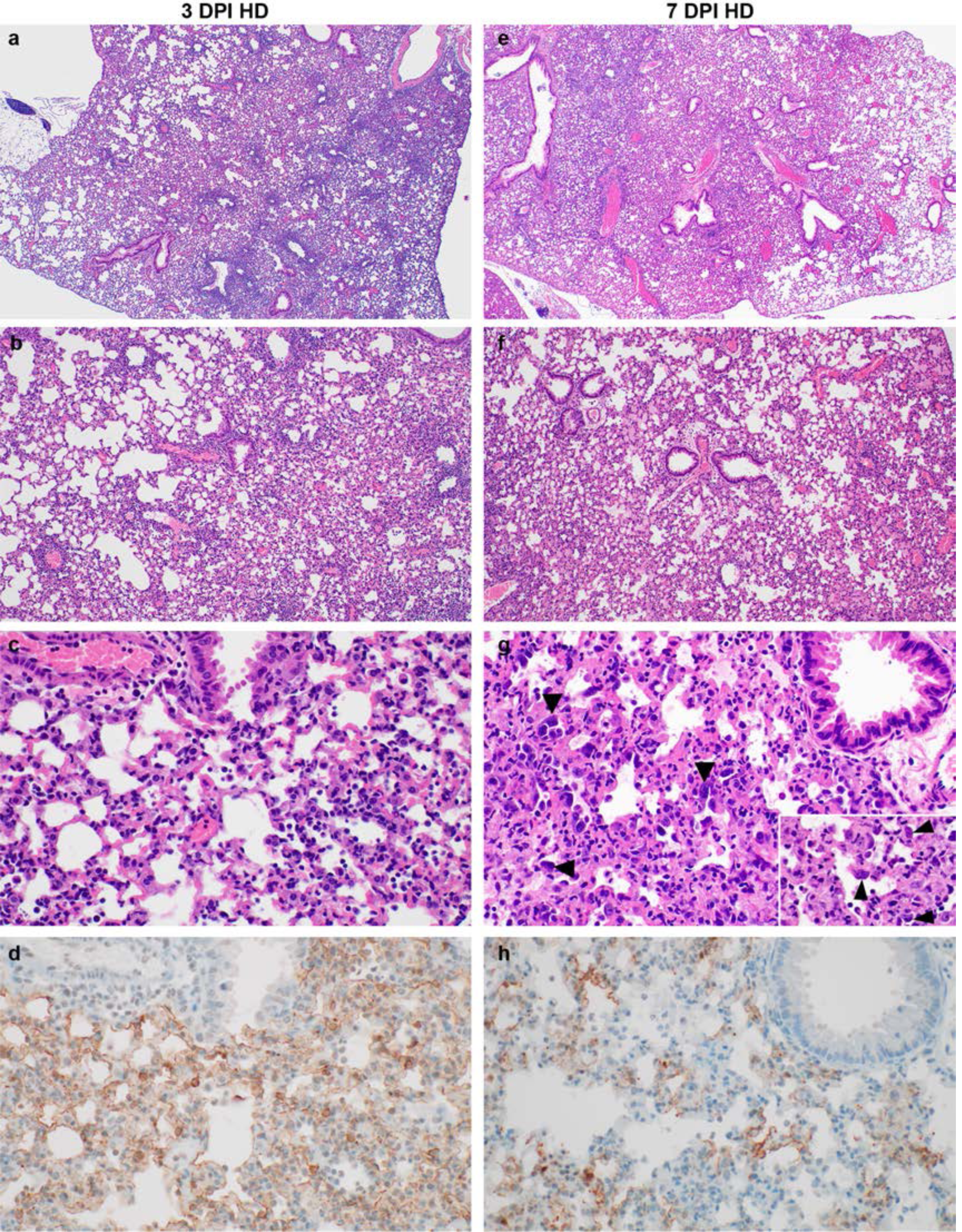
Pathological changes in lungs of K18-hACE mice inoculated with SARS-CoV-2 at 3 and 7 DPI. **a, b, c**. Interstitial pneumonia at 3 DPI, characterized by perivascular and septal inflammation with neutrophils, macrophages, lymphocytes, and edema. **d**. SARS-CoV-2 antigen immunoreactivity at 3 DPI in alveolar pneumocytes and macrophages. **e, f, g.** Multifocal interstitial pneumonia at 7 DPI, characterized by type II pneumocyte hyperplasia (arrowheads), alveolar and perivascular inflammation, fibrin, edema, syncytial cells (insert arrowheads), and single cell necrosis. **h**. SARS-CoV-2 antigen immunoreactivity in pneumocytes and macrophages at 7 DPI. HD: high dose (10^5^ TCID_50_ SARS-CoV-2). Magnification: a, e = 40 x; b, f = 100 x; c, g,h = 400 x, inset 1000 x.

We evaluated the localized infiltration of innate and adaptive immune cell populations at 3 and 7 DPI, as compared to control animals and the survivor at 21 DPI. An absence of immunoreactive macrophages (CD68+) in the γ-irradiated SARS-CoV-2 inoculated controls was noted (Fig 4a). In contrast, in lung tissue of infected animals, an infiltration of a limited number of macrophages at 3 and 7 DPI was seen, which persisted in the survivor up until 21 DPI (Fig 4 d, g and j). We next assessed lymphocyte infiltration into the lung in more detail. T cells were present in low numbers in the non-infected control (Fig 4b). At 3 DPI T cells numbers increases in perivascular tissue and alveolar septa and persisted through 7 DPI. B cells were present in low numbers in the γ-irradiated SARS-CoV-2 inoculated controls and at 3 DPI, increased numbers were observed in alveolar septa. B cells persisted through 7 DPI, when they started to cluster and form aggregates. At 21 DPI, T cells were found throughout the whole lung section and formation of lymphoid aggregates with B cells in perivascular tissues was observed in the survivor (Fig 4e, h and k). Interestingly, this animal also still demonstrated mildly inflamed alveolar septa which were often accompanied by foamy macrophages within affected alveoli (S2 Fig).

**Fig 4.**
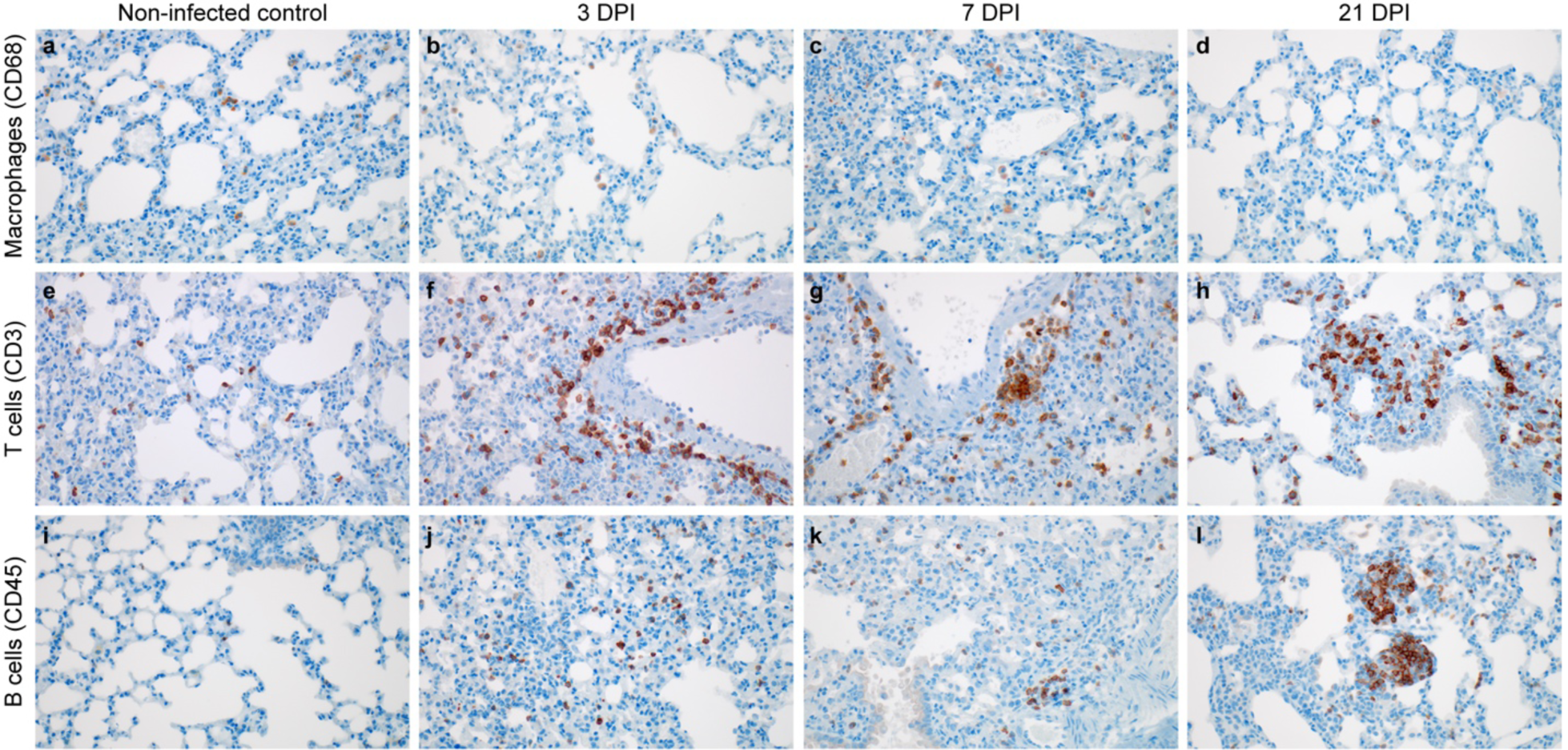
Infiltration of innate and adaptive immune-cell populations in the lungs of SARS-CoV-2 infected mice. **a-c. a, e, i**. γ-irradiated SARS-CoV-2 inoculated controls. **b, f, j**. 10^5^ TCID_50_ 3 DPI. **c, g, k**, 10^5^ TCID_50_ 7 PDI. **d, h, l**. survivor animal 21 DPI. **a.** Controls (animals inoculated with γ-irradiated SARS-CoV-2) with few macrophages (brown). **b, c.** Increased macrophages (brown) at 3 and 7 DPI. **d.** Macrophages (brown) present at end of study in a surviving mouse**. e.** Scattered T cells (brown) in the non infected control. **f, g.** T cells (brown) are increased in perivascular tissue and alveolar septa at 3 and 7 DPI. **h**. T cells (brown) forming lymphoid aggregates with B cells in perivascular tissues. **i.** B cells (brown) are few in the non infected control. **j and k.** B cells (brown) are increased in alveolar septa at 3 and 7 DPI. **l.** B cells (brown) forming lymphoid aggregates with T cells in perivascular tissues. Magnification: a-l = 400 x.

Both SARS-CoV-2 inoculated groups showed only limited lesions in the nasal turbinates at 3 and 7 DPI (Fig 5a-5b). IHC showed multifocal SARS-CoV-2 antigen in ciliated respiratory epithelial cells (Fig 5c-5d).

**Fig 5.**
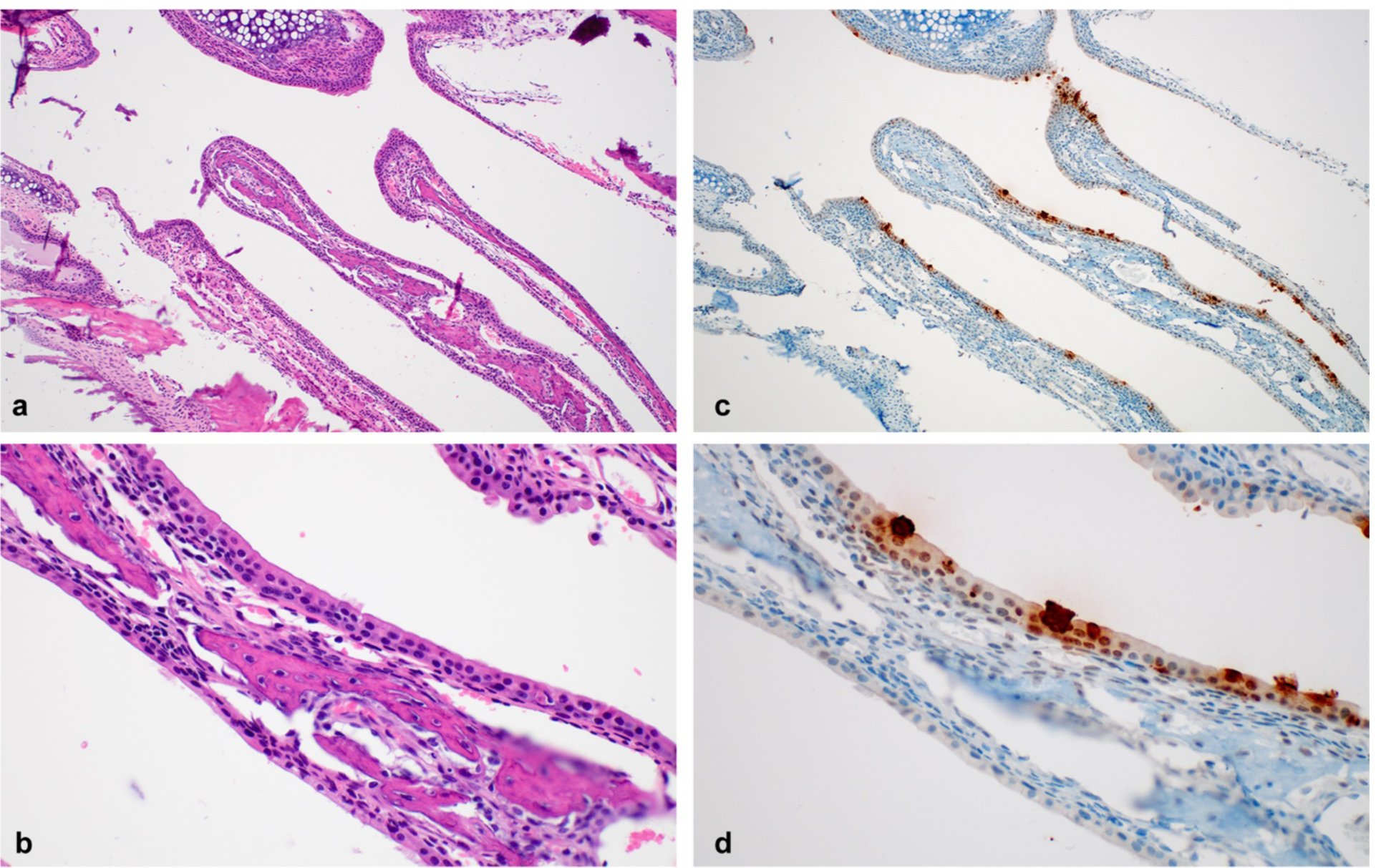
Pathological changes in nasal turbinates of SARS-CoV-2 infected mice. **a**. Nasal turbinates lined by respiratory epithelium **b**. SARS-CoV-2 antigen (brown) visible in respiratory epithelial cells. **c.** nasal turbinates without inflammation. **d.** Viral antigen in the cytoplasm of ciliated respiratory epithelial cells. Magnification: a, c = 100 x; b, d = 400 x.

At 3 DPI all brains were histologically normal (Fig 6a-6b). However, 7 DPI brain tissues showed lesions raging from minimal to moderate and included lymphocytic perivascular cuffing, gliosis, meningitis, encephalitis and microthrombi, a generalized increase in cellularity of the meninges, cerebral cortex and hippocampus and presence of edema (Fig 6c-6d). Abundant SARS-CoV-2 antigen was detected in the cerebral cortex and hippocampus within neurons and glial cells along the soma and axons (Fig 6e-6f). In addition, cerebral cortex contained microthrombi and an increased glial cell count, infiltration of inflammatory cells and scant hemorrhage (Fig 6g-6f).

**Fig 6.**
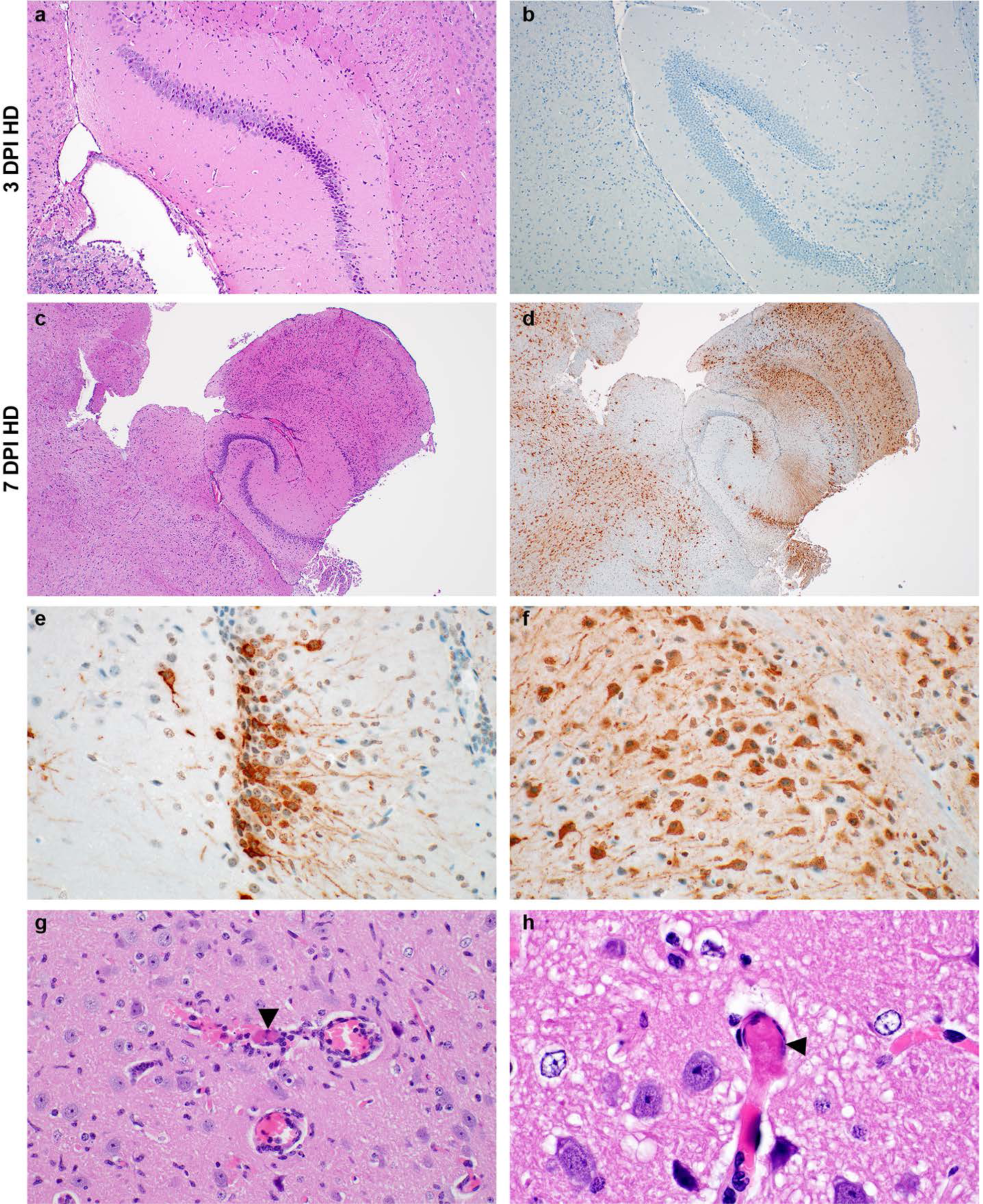
Neurotropism of SARS-CoV-2 in infected mice at 7 DPI. **a and b**. Normal hippocampus with no SARS-CoV-2 antigen detected at 3 DPI. **c**. Generalized increase in cellularity of the cerebral cortex and hippocampus; meninges are mildly expanded by edema and inflammatory cells at 7 DPI. **d**. SARS-CoV-2 antigen (brown) visible throughout the cerebral cortex and hippocampus at 7 DPI. **e and f**. SARS-CoV-2 antigen in neurons of the hippocampus and cerebral cortex highlights the soma and axons at 7 DPI. **g**. A small caliber vessel in the cerebral cortex contains a microthrombus (arrowheads) surrounded by hemorrhage and inflammatory cells which infiltrate the adjacent neuropil; there are increased glial cells throughout the image. **h**. Another microthrombus (arrowheads) in a small caliber vessel. a-b = 3 DPI, c-h = 7 DPI, dose group = 10^5^ TCID_50_ SARS-CoV-2. Magnification: a, b = 100 x; c, d = 40 x; e-g 400 x; and h = 1000 x.

### Rapid humoral immune response in SARS-CoV-2-inoculated K18-hACE mice

We next investigated two key aspects of the anti-viral immune response. To assess B-cell response and class-switch, the presence of SARS-CoV-2 spike-specific immunoglobulin (Ig)G and IgM antibodies in serum obtained at 3 and 7 DPI was investigated using ELISA. By 3 DPI, one mouse in the high dose group was positive for IgM and no mice were positive for IgG. In contrast, both spike-specific IgM and IgG were found in sera of all mice at 7 DPI (Fig 7a). IgM and IgG titers of one surviving animal at 21 DPI were comparable to those at 7 DPI.

**Fig 7.**
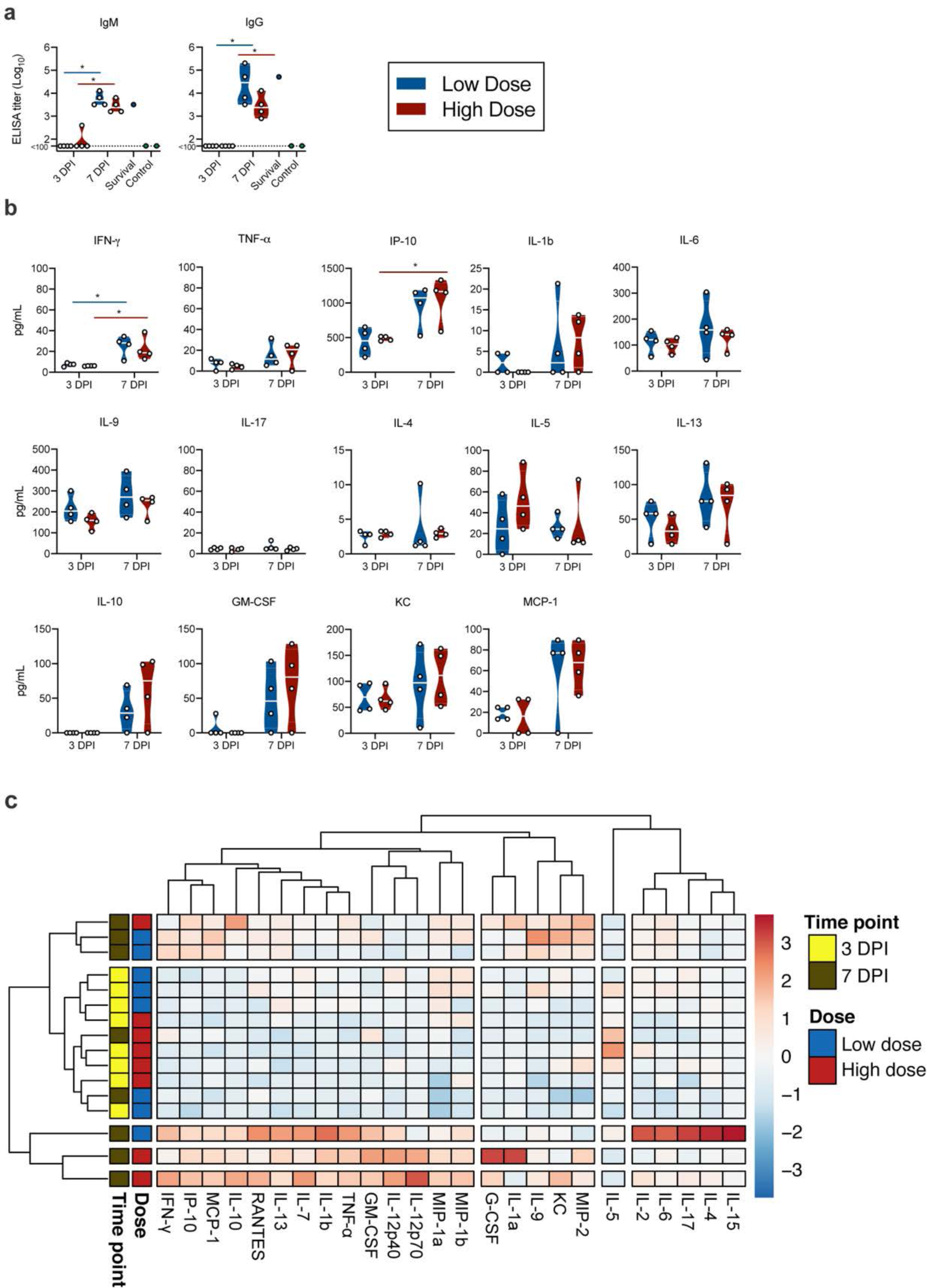
Humoral and cytokine/chemokine responses to SARS-CoV-2 infection in K18-hACE mice. **a**. IgM and IgG antibody titres against SARS-CoV-2 spike ectodomain by ELISA in serum. White line represents geometric mean of end point dilutions per study group. Dotted line represents limit of detection. **b**. Four-fold serial-diluted serum of selected cytokines/chemokines in K18-hACE mice challenged with SARS-CoV-2 measured on Bio-Plex 200 instrument (Bio-Rad) using Milliplex Mouse Cytokine/Chemokine MAGNETIC BEAD Premixed 25 Plex Kit (Millipore). Whitened represent geometric mean of all mice. **c**. Heatmap showing cytokine titers clusters based on DPI and dose of inoculation.

### Rapid systemic upregulation of proinflammatory cytokines and chemokines in SARS-CoV-2-inoculated K18-hACE mice

To investigate the immune response further we utilized serum multiplex cytokine analysis to characterize the inflammatory status and identify key patterns. Interestingly, while serum cytokine levels at 3 DPI showed only slight changes as compared to control animals, strong upregulation was observed for multiple cytokines and chemokines by 7 DPI (Fig 7b). A strong increase in T helper (Th)1-mediated cytokines interferon (IFN)-γ (both doses, p = 0.0268, 0.0268) and tumour necrosis factor (TNF)-α, (though not statistically significant) was observed. In addition, there was also an upregulation of proinflammatory and chemoattractant cytokine IFN-γ-induced protein (IP)-10 (C-X-C motif chemokine ligand (CXCL10)) (high dose, p = 0.0268). Interestingly, no trend of upregulation of Th2 anti-inflammatory cytokines interleukin (IL)-4 and IL-5 was seen, but increased levels of IL-10 were observed at DPI 7 in both groups, which has been shown to have an anti-inflammatory regulatory function in mediating antiviral responses (24). In addition, granulocyte-macrophage colony-stimulating factor (GM-CSF), KC (CXCL1) and monocyte chemoattractant protein-1 (MCP-1 (C-C motif chemokine ligand (CCL1)) were detected systemically and at increased levels at 7 DPI, further indicating a systemic recruitment of inflammatory and innate immune cells to sites of infection (Fig 3a and S2a Fig). Of note, this model did not recapitulate the increase of systemic IL-6 observed in severe COVID-19 patients (25) in either dose or timepoint. When comparing the overall cytokine profile of each animal, it became obvious that there was a stronger link between time post inoculation than between the viral dose and the resulting cytokine upregulation. We observed 3 clusters, which showed a clear time-correlation and did not detect significant differences between low and high dose inoculated animals (Fig 7c). Correlation of serum cytokine expression with lung viral gRNA did not reveal any significant positive correlation (S2b Fig).

## Discussion

In humans, COVID-19 has a broad clinical spectrum ranging from asymptomatic to severe disease (4-6, 25). Wildtype mice are not susceptible to infection with SARS-CoV-2 due to an inability of mACE2 to facilitate sufficient cellular entry (17, 18). Based on existing lethal mouse models for SARS-CoV, first described by McCray and colleagues (20), several transgenic mouse models for COVID-19 have been developed using expression of hACE2 (21, 26-29). However, mice expressing hACE2 under the mACE2 promoter (21, 26) or exogenously transfected with hACE2 showed only moderate disease with slight weight loss, reduced lung pathology and no lethal phenotype (27, 29). A mouse model expressing hACE2 under a lung ciliated epithelial cell HFH4 promoter exhibited generally only mild symptoms with lethality observed only in animals with brain infection (28). In contrast, the K18-hACE2 mouse model described here, which expresses hACE2 under the K18 epithelial promotor, displayed a high morbidity and mortality in both high dose and low dose groups. These findings are corroborated by two other studies, currently in preprint (30, 31), which demonstrate a similar disease phenotype in this model.

Previous experiments in different hACE2 mice have demonstrated varying degrees of lung pathology upon infection with SARS-CoV-2 (19, 21-23). The K18-hACE2 mice developed edema-associated acute lung injury similar to the clinical features of COVID-19 patients, including histological aspects of ARDS. This is in line with observations made in HFH4-hACE2 mice and mice expressing hACE2 under control of the murine ACE2 promotor, where viral RNA was also detected in brain tissues (28). Severe COVID-19 is histologically characterized by diffuse alveolar damage with hyaline membranes, edema, fibrin deposits, multinucleated cells, type II pneumocyte hyperplasia and lymphocyte infiltration composed of a mixture of CD4 and CD8 lymphocytes (32-34). The analyses of the pathological response observed within the lungs of the SARS-CoV-2 infected mice resemble those observed in humans with regards to lesions and cell tropism.

In humans, systemic cytokine response to SARS-CoV-2 infection are comprised of TNF-α, IL-1β, IL-1Rα, sIL-2Rα, IL-6, IL-10, IL-17, IL-18, IFN-γ, MCP-3, M-CSF, MIP-1α, G-CSF, IP-10, and MCP-1 (35-37). In the lungs of aged hACE2 mice, SARS-CoV-2 infection leads to elevated cytokine production including Eotaxin, G-CSF, IFN-γ, IL-9, and MIP-1β (38). Here, we show that SARS-CoV-2 infection of K18-hACE2 mice elicits a measurable systemic pro-inflammatory cytokine response which is significantly increased at 7 DPI and characterized by an increase in IFN-γ, TNF-α and IP-10, and also encompasses upregulation of innate cell-recruiting chemokines GM-CSF and MCP-1. Importantly, increased levels of IFN-γ, IP-10, MCP-1 and TNF-α are associated with severity of disease in in COVID-19 patients (35, 39, 40). COVID-19 patients also show heightened IL-4 and IL-10 levels, cytokines associated with inhibitory inflammatory responses (41). While the K18-hACE2 model did not recapitulate IL-4 upregulation, increased IL-10 levels were observed in serum, suggesting that both pro- and anti-inflammatory cytokine response are functioning in this mouse model. This is particularly relevant, as in COVID-19, the resulting cytokine storm is not only thought to be detrimental to disease progression but also closely linked to the development of ARDS (39). In addition, cytokine levels are also reported to be indicative of extrapulmonary multiple-organ failure (42, 43). Reports suggest that upregulation of IL-6, IL-8, and TNF-α contributes to SARS-related ARDS (35, 44). Interestingly, while we did observe the upregulation of TNF-α, IL-6 levels remained unchanged. This needs to be further investigated to clarify if our observation suggests a differently modulated immune response and pathogenesis that should be considered for intervention studies.

We have also demonstrated a functional humoral immune response and production of both IgM and IgG antibodies. This is in line with observations made in ACE2-HB-01 mice where IgG antibodies against spike protein of SARS-CoV-2 were also observed (26). This indicates that the K18-hACE2 mouse model mounts a robust innate and adaptive immune response.

The mouse model presented here recapitulates histopathological findings of COVID-19 associated ARDS, a robust innate and adaptive immune-response, neurological involvement and, importantly, presents a dose-dependent sub-lethal disease manifestation. As such, we believe this model to be highly suitable for testing of SARS-CoV-2 countermeasures such as antiviral and immune-modulatory interventions. However, COVID-19 associated ARDS in patients presents not just with characteristic lung pathology, but also with clinical manifestations including hypoxia, loss of lung compliance and requirement for intubation, liver and kidney involvement and associated increase in serum protein levels, and decreased lymphocyte numbers. To accurately assess how well K18-hACE2 mice recapitulates human ARDS, additional studies specifically addressing these aspects are required.

## Materials and Methods

### Ethics Statement

Animal experiment approval was provided by the Institutional Animal Care and Use Committee (IACUC) at Rocky Mountain Laboratories. Animal experiments were executed in an Association for Assessment and Accreditation of Laboratory Animal Care (AALAC)-approved facility by certified staff, following the basic principles and guidelines in the NIH Guide for the Care and Use of Laboratory Animals, the Animal Welfare Act, United States Department of Agriculture and the United States Public Health Service Policy on Humane Care and Use of Laboratory Animals. The Institutional Biosafety Committee (IBC) approved work with infectious SARS-CoV-2 virus strains under BSL3 conditions. All sample inactivation was performed according to IBC approved standard operating procedures for removal of specimens from high containment.

### Cells and virus

SARS-CoV-2 strain nCoV-WA1-2020 (MN985325.1) was provided by CDC, Atlanta, USA. Virus propagation was performed in VeroE6 cells in DMEM supplemented with 2% fetal bovine serum, 1 mM L-glutamine, 50 U/mL penicillin and 50 μg/mL streptomycin. VeroE6 cells were maintained in DMEM supplemented with 10% fetal bovine serum, 1 mM L-glutamine, 50 U/mL penicillin and 50 μg/mL streptomycin.

### Animal experiments

Four to six week-old male and female (15 animals each) transgenic K18-hACE2 mice expressing hACE2 (Jackson laboratories, USA, (20)) were inoculated intranasally (I.N.) with 25 µL sterile Dulbecco’s Modified Eagle Medium (DMEM) containing either 10^4^ TCID_50_ (low dose group, n = 14), 10^5^ TCID_50_ (high dose group, n = 14) or 10^5^ TCID_50_ γ-irradiate (45) (control group, n = 2) SARS-CoV-2. At 3 and 7 DPI, four mice from the low dose and high dose groups were euthanized, respectively, and tissues were collected. The remaining mice were utilized for end-point data collection and survival assessment. Mice were weighed and nasal, oropharyngeal and rectal swabs were taken daily. Mice were observed for survival up to 21 DPI or until they reached end-point criteria. End-point criteria included several parameters of severe disease (increased respiratory rate, hunched posture, ruffled fur and lethargy).

### RNA extraction and quantitative reverse-transcription polymerase chain reaction

Samples were collected with prewetted swabs in 1 mL of DMEM supplemented with 100 U/mL penicillin and 100 μg/mL streptomycin. Then, 140 µL was utilized for RNA extraction using the QIAamp Viral RNA Kit (Qiagen) using QIAcube HT automated system (Qiagen) according to the manufacturer’s instructions with an elution volume of 150 µL. Tissues (up to 30 mg) were homogenized in RLT buffer and RNA was extracted using the RNeasy kit (Qiagen) according to the manufacturer’s instructions. Viral RNA was detected by qRT-PCR (46). Five μL RNA was tested with the Rotor-GeneTM probe kit (Qiagen) according to instructions of the manufacturer. Ten-fold dilutions of SARS-CoV-2 standards with known copy numbers were used to construct a standard curve.

### SARS-CoV-2 spike glycoprotein enzyme-linked immunosorbent assay (ELISA)

Maxisorp plates (Nunc) were coated with 50 ng spike protein per well and incubated overnight at 4°C. After blocking with casein in phosphate buffered saline (PBS) (ThermoFisher) for 1 h at room temperature (RT), serially diluted 2-fold serum samples (duplicate, in casein) were incubated for 1 h at RT. Spike-specific antibodies were detected with goat anti-mouse IgM or IgG Fc (horseradish peroxidase (HRP)-conjugated, Abcam) for 1 h at RT and visualized with KPL TMB 2-component peroxidase substrate kit (SeraCare, 5120-0047). The reaction was stopped with KPL stop solution (Seracare) and read at 450 nm. Plates were washed 3x with PBS-T (0.1% Tween) in between steps. The threshold for positivity was calculated as the average plus 3x the standard deviation of negative control mouse sera.

### Measurement of cytokines and chemokines

Serum samples were inactivated with γ-irradiation (2 mRad) and cytokine concentrations were determined on a Bio-Plex 200 instrument (Bio-Rad) using Milliplex Mouse Cytokine/Chemokine MAGNETIC BEAD Premixed 25 Plex Kit (Millipore), according to the manufacturer’s instructions. Samples were pre-diluted 1:3 in the kit serum matrix (v:v). Concentrations below the limit of detections were set to zero. Heatmap and correlation graphs were made in R (47) using pheatmap (48) and corrplot (49) packages.

### Histology and immunohistochemistry

Necropsies and tissue sampling were performed according to IBC-approved protocols. Harvested tissues were fixed for eight days in 10% neutral-buffered formalin, embedded in paraffin, processed using a VIP-6 Tissue Tek (Sakura Finetek, USA) tissue processor, and embedded in Ultraffin paraffin polymer (Cancer Diagnostics, Durham, NC). Samples were sectioned at 5 µm, and resulting slides were stained with hematoxylin and eosin. Specific anti-CoV immunoreactivity was detected using an in-house SARS-CoV-2 nucleocapsid protein rabbit antibody at a 1:1000 dilution. Macrophage (CD68) and T-cell (CD3) immunoreactivities were detected using CD68 rabbit polyclonal antibody (Abcam) at a 1:250 dilution and prediluted CD3 rabbit monoclonal antibody (2GV6, Roche Tissue Diagnostics), respectively. For both CD68 and CD3, ImmPRESS-VR Horse anti-rabbit polymer was used as the secondary antibody (Vector Laboratories). B-cell (CD45) immunoreactivity was detected using anti CD45R rat monoclonal antibody (Abcam) at a 1:500 dilution and ImmPRESS goat anti-rat polymer (Vector Laboratories) as secondary antibody. The immunohistochemistry (IHC) assay was carried out on a Discovery ULTRA automated staining instrument (Roche Tissue Diagnostics) with a Discovery ChromoMap DAB (Ventana Medical Systems) kit. All tissue slides were evaluated by a board-certified veterinary anatomic pathologist blinded to study group allocations.

### Statistical analyses

Two-tailed Mann-Whitney’s rank tests and Wilcoxon matched-pairs rank test were conducted to compare differences between groups.

## Acknowledgements

The authors would like to thank Nathalie Thornburg and Susan Gerber for sharing of the SARS-CoV-2 isolate, Kizzmekia Corbett and Barney Graham for the plasmid encoding the full-length SARS-CoV-2 spike and Anita Mora for assistance with the Figs. This work was supported by the Intramural Research Program of the National Institute of Allergy and Infectious Diseases (NIAID), National Institutes of Health (NIH) (1ZIAAI001179-01).

## Supplementary Figure legends

**S1 Fig.**
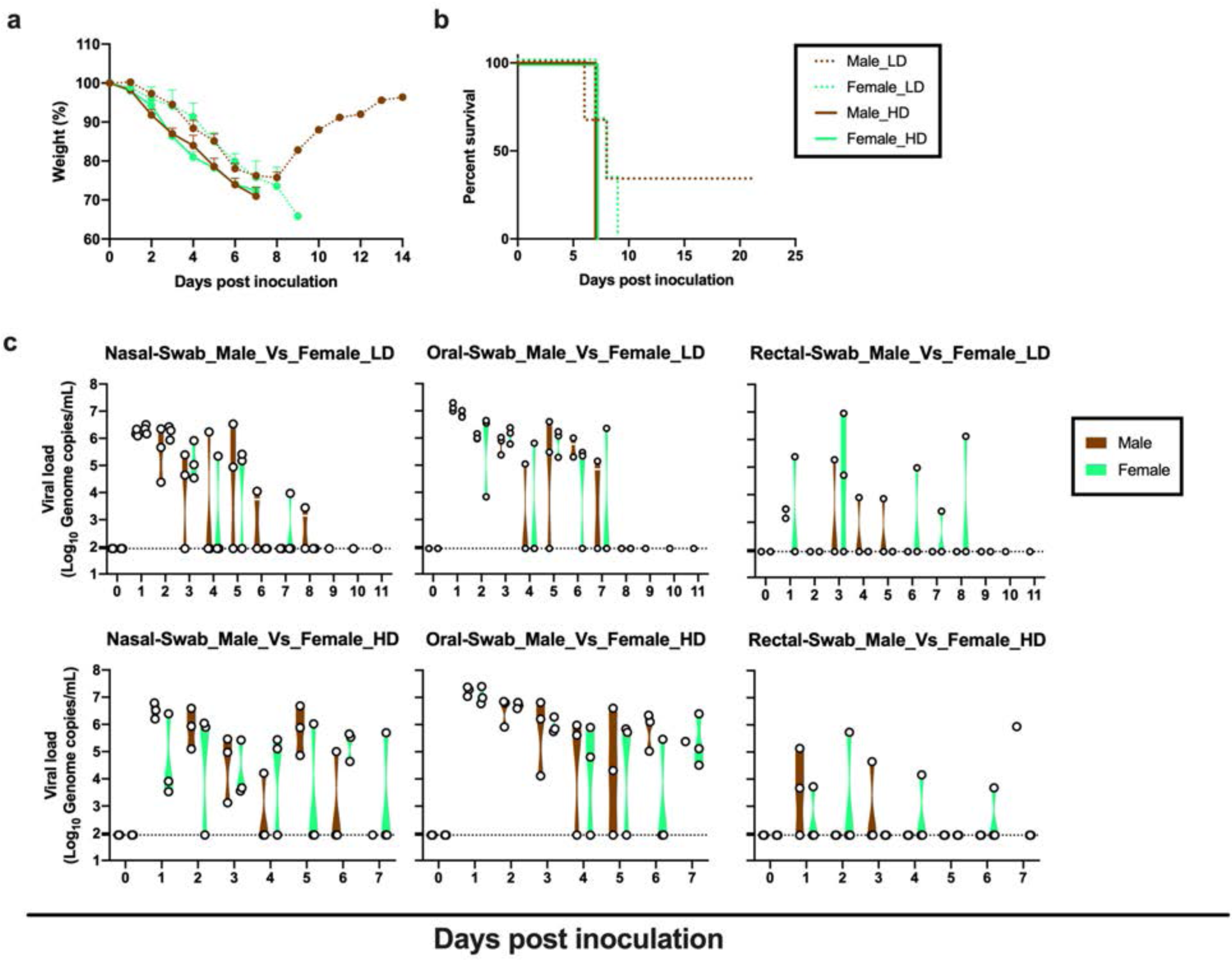
Sex-dependent weight loss, mortality and virus shedding in K18-hACE2 mice after SARS-CoV-2 infection. **a.** Body weights were monitored every day. Relative body weight changes are show for female (turquoise) and male (brown) animals for HD (solid) and LD (dotted) groups. **b.** Survival is show for female (turquoise) and male (brown) animals for HD (solid) and LD (dotted) groups. **c**. Nasal, oral and rectal virus shedding in low and high dose infected female (turquoise) and male (brown) mice was quantified by RT-qPCR across time. Individual animals are plotted, violin plot depict median and quantiles. Abbreviations: LD = low dose (10^4^ TCID_50_ SARS-CoV-2), HD = high dose (10^5^ TCID_50_ SARS-CoV-2).

**S2 Fig.**
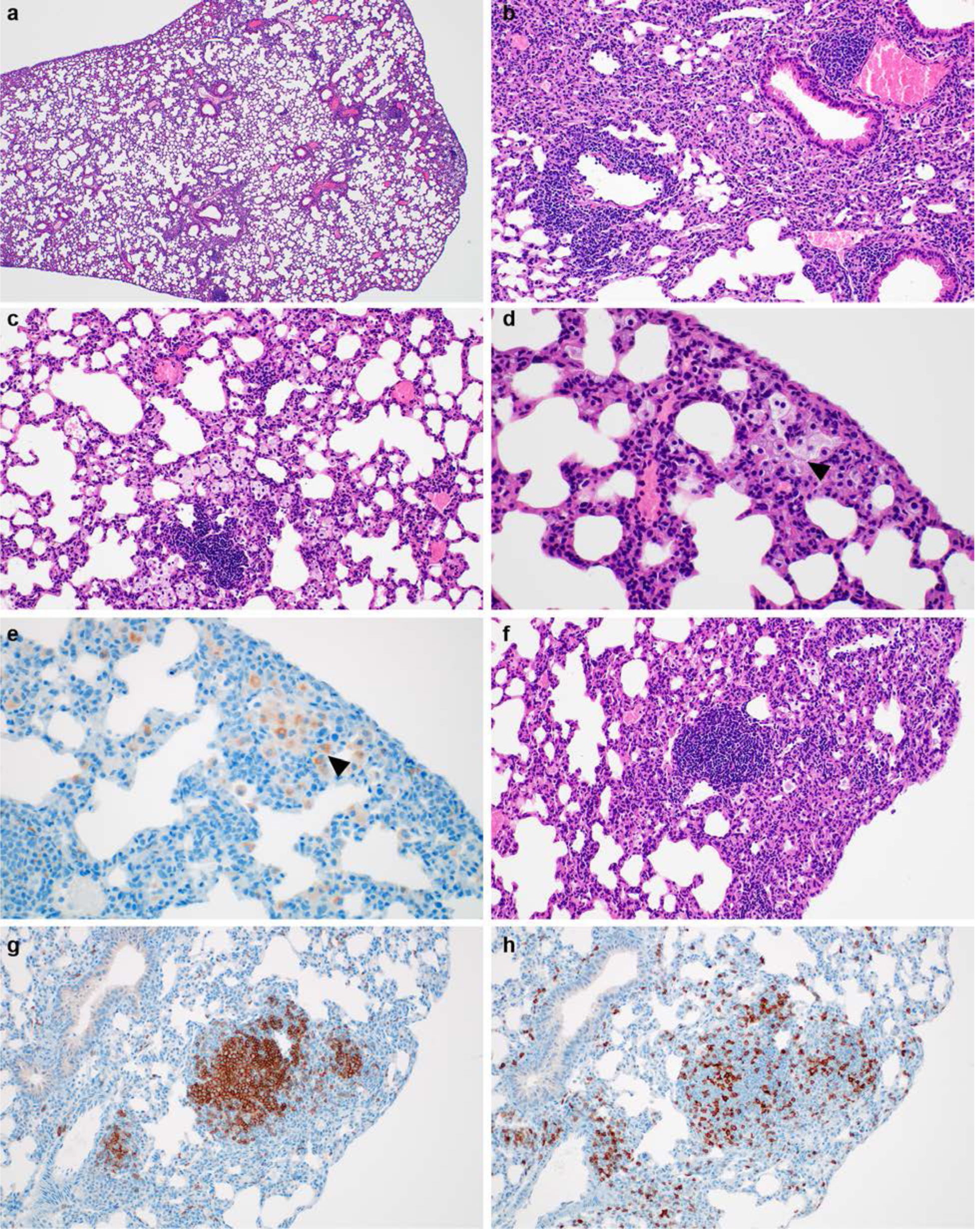
Histological analysis of lung sections from one low dose survivor at 21 days post infection. **a**. Multiple foci of perivascular inflammation and increased alveolar cellularity. **b**. Perivascular and peribronchiolar lymphocytic inflammation. **c**. Aggregated lymphocytes within alveolar septa and alveoli containing foamy macrophages. **d**. Foamy macrophages cluster and fill alveoli (arrowheads) and alveolar septa contain increased numbers of lymphocytes. **e**. CD68 immunoreactivity in foamy alveolar macrophages (arrowheads). **f**. One of many discreet aggregates of lymphocytes in the 21 DPI lung composed of **g**. CD45+ B cells and **h**. CD3+ T cells. Magnification: a = 40 x; b, c, f, g, h = 200 x; d, e = 400 x.

**S3 Fig.**
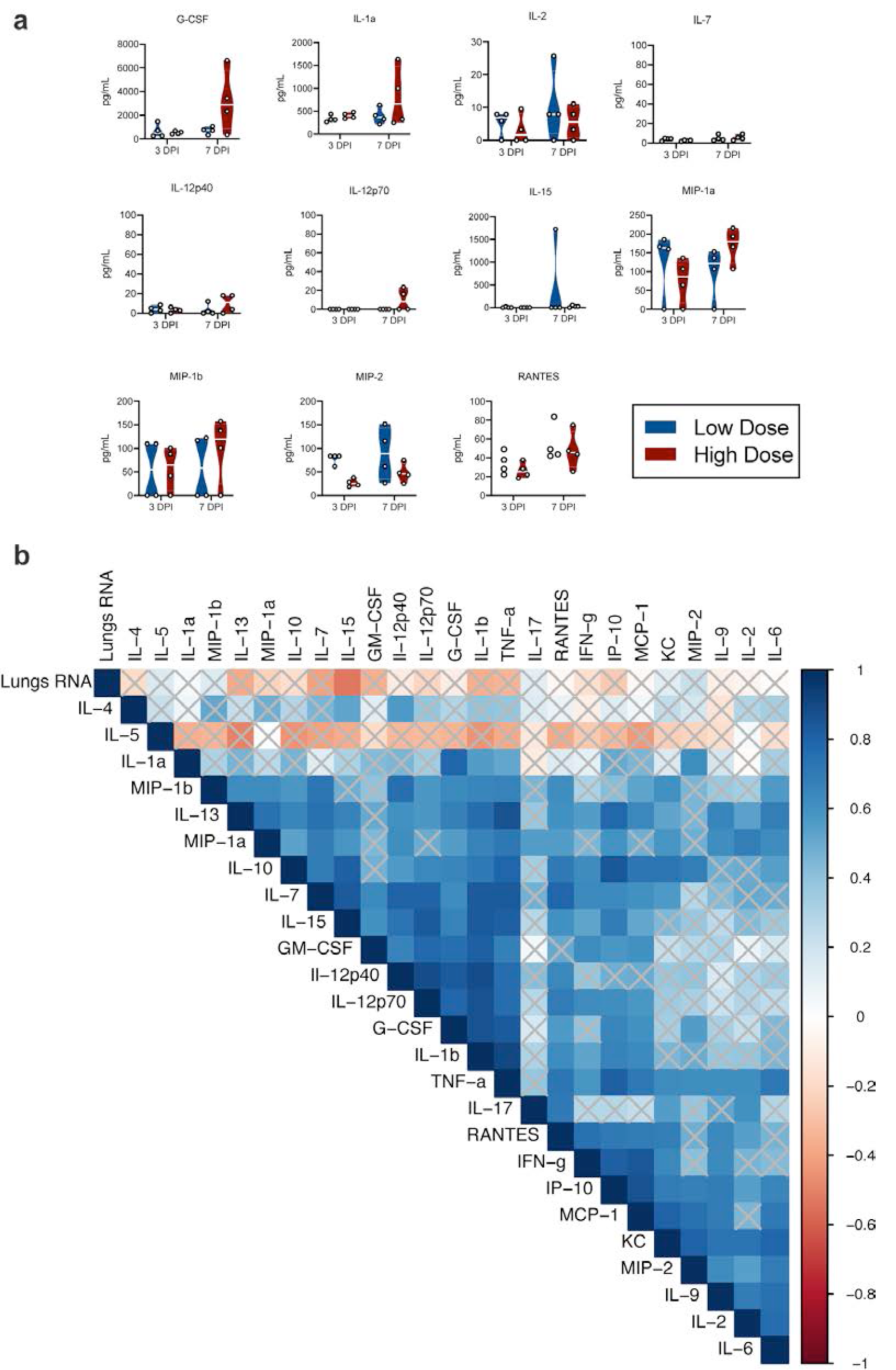
Multiplex analysis of cytokines/chemokines in K18-hACE mice challenged with SARS-CoV-2 measured at 3- and 7-days post inoculation. **a**. Individual animals are plotted, violin plots depict median and quantiles. Low dose = blue, high dose = red. **b**. Correlation between cytokine levels and viral RNA in the lungs. Significant correlations (p = 0.05) are shown and strength of correlation is depicted according to the colour bar, crossed bars are not significant. Abbreviations: DPI = days post inoculation, G-CSF = granulocyte colony-stimulating factor, GM-CSF = granulocyte-macrophage colony-stimulating factor, INF = interferon, IL = interleukin, KC = keratinocyte chemoattractant, MCP = monocyte chemoattractant protein, MIP = macrophage inflammatory protein, IP = interferon-γ-inducible protein, TNF = tumour necrosis factor.

## References

1. WHO. Coronavirus disease 2019 (COVID-19) Situation Report – 52. 2020 12 March 2020.

2. Nie S, Han S, Ouyang H, Zhang Z. Coronavirus Disease 2019-related dyspnea cases difficult to interpret using chest computed tomography. Respir Med. 2020;167:105951-.

3. Parry AH, Wani AH, Yaseen M, Dar KA, Choh NA, Khan NA, et al. Spectrum of chest computed tomographic (CT) findings in coronavirus disease-19 (COVID-19) patients in India. Eur J Radiol. 2020;129:109147-.

4. Li X, Ma X. Acute respiratory failure in COVID-19: is it “typical” ARDS? Crit Care. 2020;24(1):198.

5. Jiang F, Deng L, Zhang L, Cai Y, Cheung CW, Xia Z. Review of the Clinical Characteristics of Coronavirus Disease 2019 (COVID-19). J Gen Intern Med. 2020;35(5):1545–9.

6. Li X, Ma X. Acute respiratory failure in COVID-19: is it “typical” ARDS? Crit Care. 2020;24(1):198-.

7. Calcagno N, Colombo E, Maranzano A, Pasquini J, Keller Sarmiento IJ, Trogu F, et al. Rising evidence for neurological involvement in COVID-19 pandemic. Neurol Sci. 2020;41(6):1339–41.

8. Dhakal BP, Sweitzer NK, Indik JH, Acharya D, William P. SARS-CoV-2 Infection and Cardiovascular Disease: COVID-19 Heart. Heart Lung Circ. 2020.

9. Mao L, Jin H, Wang M, Hu Y, Chen S, He Q, et al. Neurologic Manifestations of Hospitalized Patients With Coronavirus Disease 2019 in Wuhan, China. JAMA Neurol. 2020.

10. Chan JF, Zhang AJ, Yuan S, Poon VK, Chan CC, Lee AC, et al. Simulation of the clinical and pathological manifestations of Coronavirus Disease 2019 (COVID-19) in golden Syrian hamster model: implications for disease pathogenesis and transmissibility. Clin Infect Dis. 2020.

11. Kim YI, Kim SG, Kim SM, Kim EH, Park SJ, Yu KM, et al. Infection and Rapid Transmission of SARS-CoV-2 in Ferrets. Cell Host Microbe. 2020;27(5):704–9 e2.

12. Munster VJ, Feldmann F, Williamson BN, van Doremalen N, Perez-Perez L, Schulz J, et al. Respiratory disease in rhesus macaques inoculated with SARS-CoV-2. Nature. 2020.

13. Woolsey C, Borisevich V, Prasad AN, Agans KN, Deer DJ, Dobias NS, et al. Establishment of an African green monkey model for COVID-19. bioRxiv. 2020.

14. Yu P, Qi F, Xu Y, Li F, Liu P, Liu J, et al. Age-related rhesus macaque models of COVID-19. Animal Model Exp Med. 2020;3(1):93–7.

15. Rockx B, Kuiken T, Herfst S, Bestebroer T, Lamers MM, Oude Munnink BB, et al. Comparative pathogenesis of COVID-19, MERS, and SARS in a nonhuman primate model. Science. 2020;368(6494):1012–5.

16. Letko M, Marzi A, Munster V. Functional assessment of cell entry and receptor usage for SARS-CoV-2 and other lineage B betacoronaviruses. Nature microbiology. 2020;5(4):562–9.

17. Zhou P, Yang XL, Wang XG, Hu B, Zhang L, Zhang W, et al. A pneumonia outbreak associated with a new coronavirus of probable bat origin. Nature. 2020;579(7798):270–3.

18. Zhao X, Chen D, Szabla R, Zheng M, Li G, Du P, et al. Broad and differential animal ACE2 receptor usage by SARS-CoV-2. bioRxiv. 2020.

19. Jing Sun, Zhuang Z, Zheng J, Li K, Wong RL-Y, Liu D, et al. Generation of a Broadly Useful Model for COVID-19 Pathogenesis, Vaccination, and Treatment. Cell. 2020.

20. McCray PB, Jr., Pewe L, Wohlford-Lenane C, Hickey M, Manzel L, Shi L, et al. Lethal infection of K18-hACE2 mice infected with severe acute respiratory syndrome coronavirus. J Virol. 2007;81(2):813–21.

21. Sun SH, Chen Q, Gu HJ, Yang G, Wang YX, Huang XY, et al. A Mouse Model of SARS-CoV-2 Infection and Pathogenesis. Cell Host Microbe. 2020.

22. Hongjing Gu, Chen Q, Yang G, He L, Fan H, Deng Y-Q, et al. Rapid adaption of SARS-CoV-2 in BALB/c mice: Novel mouse model for vaccine efficacy. BioRxiv. May 2, 2020.

23. Dinnon KH, Leist SR, Schafer A, Edwards CE, Martinez DR, Montgomery SA, et al. A mouse-adapted SARS-CoV-2 model for the evaluation of COVID-19 medical countermeasures. bioRxiv. 2020.

24. Rojas JM, Avia M, Martín V, Sevilla N. IL-10: A Multifunctional Cytokine in Viral Infections. J Immunol Res. 2017;2017:6104054-.

25. Chen G, Wu D, Guo W, Cao Y, Huang D, Wang H, et al. Clinical and immunological features of severe and moderate coronavirus disease 2019. J Clin Invest. 2020;130(5):2620–9.

26. Bao L, Deng W, Huang B, Gao H, Liu J, Ren L, et al. The pathogenicity of SARS-CoV-2 in hACE2 transgenic mice. Nature. 2020.

27. Hassan AO, Case JB, Winkler ES, Thackray LB, Kafai NM, Bailey AL, et al. A SARS-CoV-2 Infection Model in Mice Demonstrates Protection by Neutralizing Antibodies. Cell. 2020.

28. Jiang RD, Liu MQ, Chen Y, Shan C, Zhou YW, Shen XR, et al. Pathogenesis of SARS-CoV-2 in Transgenic Mice Expressing Human Angiotensin-Converting Enzyme 2. Cell. 2020.

29. Israelow B, Song E, Mao T, Lu P, Meir A, Liu F, et al. Mouse model of SARS-CoV-2 reveals inflammatory role of type I interferon signaling. bioRxiv. 2020:2020.05.27.118893.

30. Golden JW, Cline CR, Zeng X, Garrison AR, Carey BD, Mucker EM, et al. Human angiotensin-converting enzyme 2 transgenic mice infected with SARS-CoV-2 develop severe and fatal respiratory disease. bioRxiv. 2020:2020.07.09.195230.

31. Moreau GB, Burgess SL, Sturek JM, Donlan AN, Petri WA, Mann BJ. Evaluation of K18-<em>hACE2</em> mice as a model of SARS-CoV-2 infection. bioRxiv. 2020:2020.06.26.171033.

32. Tian S, Hu W, Niu L, Liu H, Xu H, Xiao S-Y. Pulmonary Pathology of Early-Phase 2019 Novel Coronavirus (COVID-19) Pneumonia in Two Patients With Lung Cancer. J Thorac Oncol. 2020;15(5):700–4.

33. Schaller T, Hirschbühl K, Burkhardt K, Braun G, Trepel M, Märkl B, et al. Postmortem Examination of Patients With COVID-19. JAMA. 2020;323(24):2518–20.

34. Fox SE, Akmatbekov A, Harbert JL, Li G, Brown JQ, Vander Heide RS. Pulmonary and Cardiac Pathology in Covid-19: The First Autopsy Series from New Orleans. medRxiv. 2020:2020.04.06.20050575.

35. Yang Y, Shen C, Li J, Yuan J, Wei J, Huang F, et al. Plasma IP-10 and MCP-3 levels are highly associated with disease severity and predict the progression of COVID-19. J Allergy Clin Immunol. 2020;146(1):119-27.e4.

36. Liu Y, Zhang C, Huang F, Yang Y, Wang F, Yuan J, et al. Elevated plasma levels of selective cytokines in COVID-19 patients reflect viral load and lung injury. National Science Review. 2020;7(6):1003–11.

37. Wan S, Yi Q, Fan S, Lv J, Zhang X, Guo L, et al. Characteristics of lymphocyte subsets and cytokines in peripheral blood of 123 hospitalized patients with 2019 novel coronavirus pneumonia (NCP). medRxiv. 2020:2020.02.10.20021832.

38. Sun S-H, Chen Q, Gu H-J, Yang G, Wang Y-X, Huang X-Y, et al. A Mouse Model of SARS-CoV-2 Infection and Pathogenesis. Cell host & microbe. 2020;28(1):124-33.e4.

39. Ye Q, Wang B, Mao J. The pathogenesis and treatment of the ‘Cytokine Storm’ in COVID-19. J Infect. 2020;80(6):607–13.

40. Huang C, Wang Y, Li X, Ren L, Zhao J, Hu Y, et al. Clinical features of patients infected with 2019 novel coronavirus in Wuhan, China. The Lancet. 2020;395(10223):497–506.

41. Chen L, Liu HG, Liu W, Liu J, Liu K, Shang J, et al. [Analysis of clinical features of 29 patients with 2019 novel coronavirus pneumonia]. Zhonghua Jie He He Hu Xi Za Zhi. 2020;43(0):E005.

42. Wang H, Ma S. The cytokine storm and factors determining the sequence and severity of organ dysfunction in multiple organ dysfunction syndrome. Am J Emerg Med. 2008;26(6):711–5.

43. Parsons PE, Eisner MD, Thompson BT, Matthay MA, Ancukiewicz M, Bernard GR, et al. Lower tidal volume ventilation and plasma cytokine markers of inflammation in patients with acute lung injury. Crit Care Med. 2005;33(1):1-6; discussion 230-2.

44. Girija ASS, Shankar EM, Larsson M. Could SARS-CoV-2-Induced Hyperinflammation Magnify the Severity of Coronavirus Disease (CoViD-19) Leading to Acute Respiratory Distress Syndrome? Frontiers in Immunology. 2020;11(1206).

45. Feldmann F, Shupert WL, Haddock E, Twardoski B, Feldmann H. Gamma Irradiation as an Effective Method for Inactivation of Emerging Viral Pathogens. The American journal of tropical medicine and hygiene. 2019;100(5):1275–7.

46. Corman VM, Landt O, Kaiser M, Molenkamp R, Meijer A, Chu DK, et al. Detection of 2019 novel coronavirus (2019-nCoV) by real-time RT-PCR. Euro Surveill. 2020;25(3).

47. R Development Core Team. R: A language and Environment for Statistical computing R Foundation for Statistical Computing; 2010.

48. Kolde R. Implementation of heatmaps that offers more control over dimensions and appearance. 2019.

49. Wei T, Simko V. R package “corrplot”: Visualization of a Correlation Matrix. Version 0.84 ed2017.

